# Simultaneous Multisheet Multichannel Imaging Cytometry (SMMIC) System Powered by Adaptive Vertically-Aligned Multi-Sheet Array (VAMSA) PSF

**DOI:** 10.1101/2022.06.25.497275

**Authors:** Prakash Joshi, Prashant Kumar, S Aravinth, Partha P. Mondal

## Abstract

Volumetric interrogating of a large population of live specimens at high throughput is a challenging task that necessitates new technology. We propose vertical-aligned multi-sheet array (VAMSA) illumination PSF that enables interrogation of specimens flowing simultaneously through multiple microfluidic channels. The very geometry of PSF enables high quality cross-sectional imaging, and facilitates volumetric interrogation of specimens flowing through commercial microfluidic chip (consists of multiple flow-channels), which is a step towards large population screening. The SMMIC technique employs a unique combination of transmission grating, beam-expander and high NA objective system in a specific optical configuration to generate diffraction-limited illumination PSF (VAMSA-PSF). However, the detection is accomplished by a large field-of-view widefield 4f-system that consists of low NA objective lens, high performance fluorescence filters, and tube lens. Studies show high quality sectional images (resolution ∼ 2.5*μm*, and SBR ∼ 4.8*dB*) of HeLa cancerous cells at high flow throughput (flow-rate of, 2500 *nl/min*). A cell count of > 1*k* and volume reconstruction efficiency of ∼ 121 *cells/min* is noteworthy. In addition, SMMIC system demonstrate organelle-level resolution with a SBR comparable to that of confocal especially at low flow-rates. It is hoped that the proposed system may accelerates drug-treatment studies for a large population of live specimens to advance the evolving field of translational medicine and health-care.

---

To be able to carry out multiple tasks (volume visualization, biophysical parameter estimation, and statistical analysis) on the go is a demanding task that potentially calls for new technological breakthrough at the fundamental level. This is something true for the field of imaging flow cytometry which is still depending on the traditional point-illumination based technology [1] [2] [3] [4] [5] [6] [7] [8]. Although existing techniques have found numerous applications, further advance is possible if new capabilities are integrated to make it a full-fledged diagnostic system. In this respect, the arrival of light sheet has shown promise and potential to advance the field to match the current expectations. The research disciplines that are most likely to get boost from the emerging light sheet technology range from translational medicine to clinical health-care.

Since its first inception in the year 2013, light sheet cytometry has been used for a variety of studies [22] [23]. In the year 2016, Lau et al reported light sheet based optofluidic time-stretch imaging for high throughput interrogation of MIHA cells [24]. An exciting development is the on-chip cytometry realized by integrating light sheet on a mirror embedded microfluidic chip [25]. Other variants include scanned Bessel beam for stem cell research, and a label-free imaging flow cytometry for cell screening [27] [26]. On similar lines, Jiang et al have realized light sheet cytometry by integrating droplet microfluidics and light sheet [29]. In another development, successful imaging of large live organisms (such as, C elegans) is achieved by iLIFE imaging cytometry [28], and SCAPE especially at high speed [30]. The developments continued, with our group succeeding in high speed interrogation of cells using a single light sheet for high-throughput imaging [31], and a line excitation array detection (LEAD) fluorescence microscopy that performs close to mega-Hertz line-scanning of an excitation laser line using a virtual light sheet [32]. Another distinct advance has been the use of high throughput imaging cytometry system for disease diagnosis [33]. There are many such advancements made in the past few years that have consolidated and brought forward the potential of light sheet powered imaging cytometry. Here, we report a full-fledged multiple light sheet imaging cytometry that can be used for a variety of studies.

The technology largely employed in present state-of-the art imaging cytometry systems are based on point-illumination that necessitates colineating cells to enable sequential interrogation. This necessitates two fluids of which one is the specimen flowing fluid and the second is sheath fluid required for hydrodynamic focusing of the flowing specimens [34] [35]. This arrangement makes the system complex and quite cumbersome to operate. The existing systems are not suitable for all kinds of specimens (such as, cells, tissues and organisms) and needs special arrangements for optimal operation. Moreover, existing point-based cytometry techniques are largely limited to counting, sorting and mixing. Although limited, some of the developments are worth mentioning. Existing techniques has found several technique in diverse research disciplines ranging from cell biology (asymmetric cell division, receptor activity, autophagy, nuclear translocation and cell cycle analysis) to medical health-care (drug-discovery, disease biology and immunology) [9] [10] [11] [12] [13] [14] [15] [16] [17] [18]. Moreover, recent advancement in complex multichannel cell analysis based on machine learning techniques have advanced point-illumination based cytometry techniques [19] [20] [21]. Apart from these benefits, there are some limitations as well. One of the key obstacle with existing techniques is their inability to images internal structure in a specimen (such as,cell) and thus are not particularly suitable for studying underlying biological mechanisms in live specimens. Specifically in drug-discovery, it would be advantageous to interrogate a large population of cells (both healthy and diseased) with the ability to visualize internal organelles and their dynamics. The compelling properties of light sheet that makes it stand out are, large field-of-view (suitable for cross-sectioning of large specimens), selective plane interrogation, and better SBR. All together, the current limitation of imaging cytometry some what limits its use to simple trivial tasks, and new technology needs to step up to further its progress. In the present scenario, light sheet technology seems to provide such a capability as evident from recent developments. A detailed discussion of light sheet technology and its applications in diverse disciplines can be found in Ref. [36] [37] [38].

In this article, we propose vertical-aligned multi-sheet array (VAMSA) illumination for imaging cytometry on a commercial *Y* -type microfluidic chips. The system is calibrated and tested on a known test sample (fluorescent beads in solution) for optimal operation. Subsequently, the system is employed to localize internal organelles (mitochondria) in cancerous HeLa cells and map its distribution. In the process, critical biophysical parameters related to cell and internal organelles dynamics are also estimated.

The schematic diagram of developed vertical light sheet based SMMIC system is shown in Fig. 1. SMMIC has three major sub-system apart from fast data acquisition system. The illumination sub-system is at the heart of SMMIC, that comprise of three major optical components arranged in a specific configuration to generate the desired illumination PSF (VAMSA). A light of wavelength 532 *nm* is used as an excitation source for the beads and fluorescently-labelled HeLa cells. The laser light is directed to a beam-stearing system (consists of a series of mirrors) for precision alignment. Subsequently the beam is passed through a specialized transmission grating beam-splitter (80 grooves / mm, Edmund Optics, Singapore) to derive a maximum of five intense beams. The beam is then subjected to the beam-expander for 2*X* expansion of the beam to fill the back-aperture of cylindrical lens. The enlarged beams then go through a combination of cylindrical and high NA objective lens to realize collinear but well-separated multiple vertical light sheets. The second sub-system consists of a fabricated microfluidic chip (containing multiple *Y*-type 200 × 100 *μm*^2^ sized flow-channels) for flowing the specimens. The illumination PSF transects the channels for optically cross-sectioning the fluorescently-labelled cells flowing through the channels. During experimentation, the cells are loaded in the reservoir and a flow-pump (New Era flow pump, NE-1002X, USA) is connected to other end for inducing flow. The pump is operated in the suction mode, and flow is computationally controlled by an interfacing software. The third sub-system is essentially an widefield 4*f* detection setup. The fluorescence from specimen (fluorescently-labelled HeLa cells) is collected by a separate objective lens (10X, 0.30 NA Olympus, Japan) and focussed by the tube lens to the detector where the image formation takes place. Note that, the objective and tube lens combination forms a 4*f* optical configuration, with a magnification of ∼ 12*X*. On its way to the detector (Zyla 4.2 sCMOS camera, Andor, UK) the light is filtered by optical filters (532 nm Notch filter, ZET 532nf, Chroma Technology, USA and 550 nm Longpass filter, Thorlabs, USA) arranged in the filter box. Finally, the data (sectional images) are recorded and cell volumes are reconstructed using the available inbuilt MATLAB scripts for data handling, deconvolution and stacking.

**FIG. 1:**
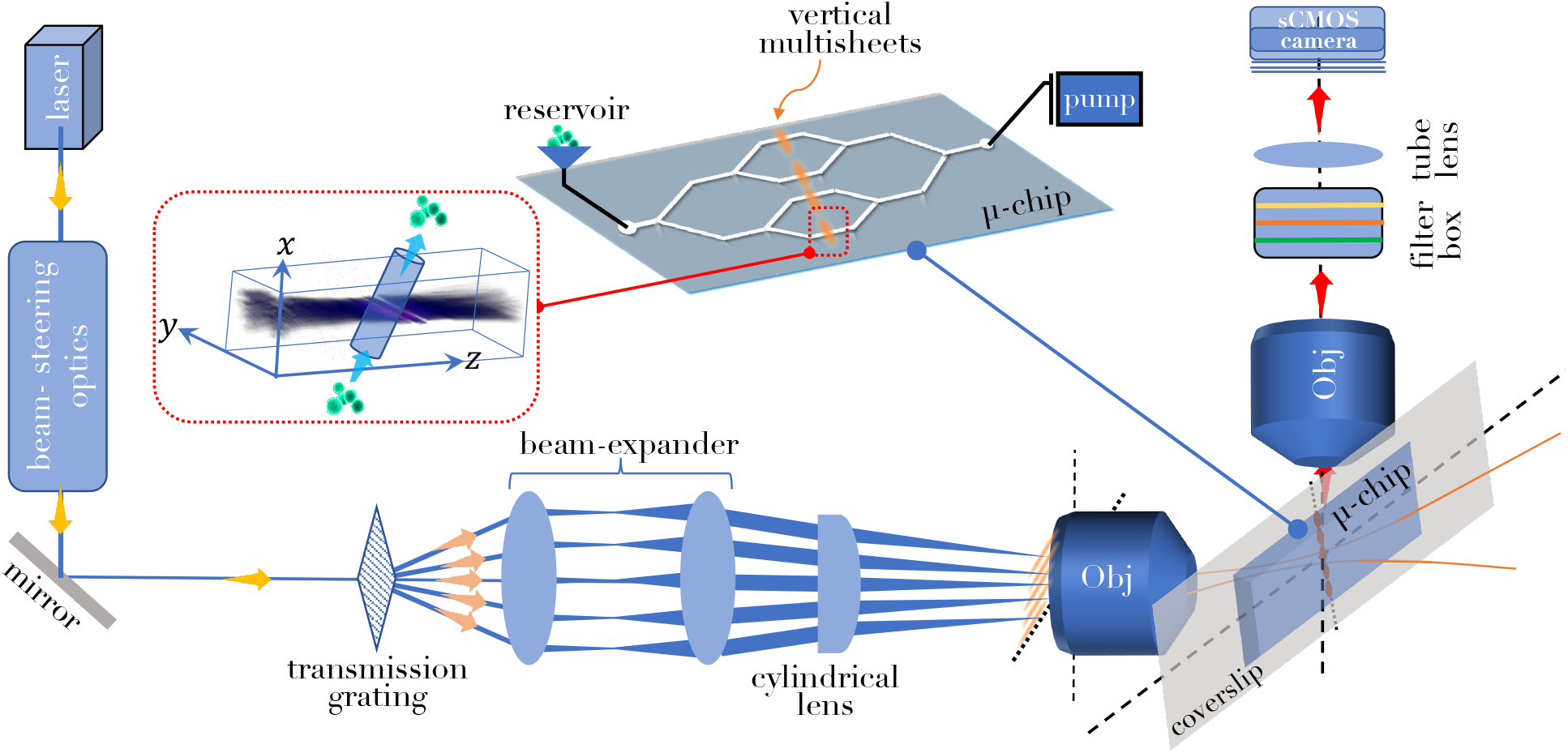
The schematic diagram of the proposed SMMIC system. The major subsystems comprise multiple vertical light sheet illumination, a microfluidic chip-based specimen flow platform, and a widefield 4*f* detection system. The illumination subsystem consists of key optical components such as transmission grating, beam-expander, cylindrical lens for realizing vertical spaced light sheets, and a high NA objective lens to generate diffraction-limited multiple vertical light sheet PSF. The inset (red box) shows a schematic realization of one of the light sheets sectioning the channel.

The next critical stage in the SMMIC system development is the characterization of illumination PSF and flow-induced detection PSF, along with other calibrations (related to flow-rate and detector exposure time). Fig. 2A shows the schematic of optical setup used for characterizing the illumination PSF. To obtain a 3D map of the illumination PSF (*h*_*ill*_), a CCD camera is placed in the beam-path of illumination sub-system, at and about the focus. The incident intensity field at every *z*-position is recorded by scanning the camera (placed on moving platform) and the images are recorded. To obtain the 3D field distribution, the images are then merged and the illumination 3D PSF is reconstructed (see, Fig. 2A, inset). An average size of the light sheet is found to be ≈ 211.0*μm* (full-width at half maximum) which can section the entire channel. Specifically, the illumination PSF shows 5 prominent peaks (corresponding to 5 channels) for a specific grating-beam-expander distance, *D* = 10 *mm*. However, the SMMIC system allows one to adaptively choosing the number of light sheets by just altering the distance between transmission grating and beam-expander. Practically, this is accomplished by placing the grating on a precision linear translator. Fig. 2B shows three, four and five light sheets for *D* = 10, 20, 25 *mm*, respectively. This brings in an additional flexibility when designing microfluidic chips with desired number of well-spaced flow channels. Since, flow induces optical aberration (blur), this has a direct effect on the recorded images. To minimize this effect, flow-variant detection PSFs are obtained by flowing pointy sources such as fluorescent beads (size 1*μm*) and recoding the same in the detection channel as shown in Fig. 2C. Calibration and analysis show a change of PSF from 4.410*μm* to 5.726*μm* with flow rate 100 to 2000 *nl/min*, respectively. Finally, the PSFs are used for deconvolving the recorded raw images before reconstructing cell volume. The deconvolution process can be calculated in the Fourier domain and expressed as, *g*(*x, y*) = *F* ^−1^[*F* (*f*)*/F* (*h*)], where, *f* (*x, y*), *g*(*x, y*) and *h*(*x, y*) represents the raw image, true image and the PSF of the detection sub-system respectively. Here, *F* (∗) represents Fourier transform.

**FIG. 2:**
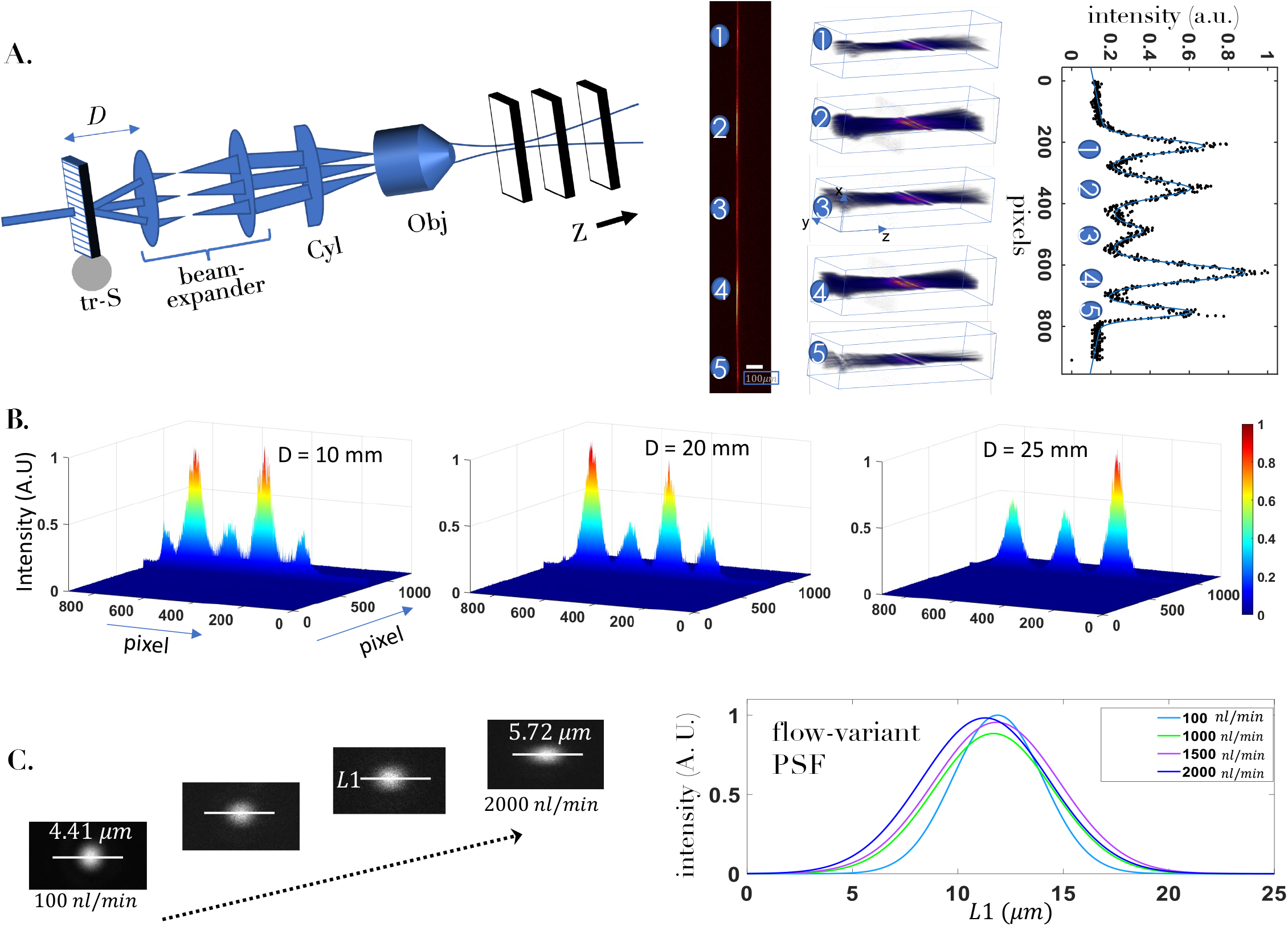
System PSF Characterization. (A) The system PSF is characterised by scanning a CCD camera along the beam path about the focus. Corresponding 2D and 3D system PSFs are also shown along with the intensity plots. (B) The change in system PSF at varying grating-beam-expander distance, *D* is also shown, suggesting adaptive e change in the number of light sheets. (C) The detection PSF at varying flow-rates from 100 *nl/min* to 2000 *nl/min*. The corresponding intensity plots shows the PSF-broadening at large flow-rates. These flow-variant PSFs are used deconvolving the recorded images.

The quantification of specimen count statistics and the reliable recording of specimen planes at specific flow-rate is critical. Our study show that at a relatively moderate flow-rate of 1000 *nl/min*, our camera can capture 5 high quality sectional images, whereas high flow-rate (> 2000 *nl/min*) result in relatively low quality images. Both beads and cell specimens are flown and the corresponding count statistics is estimated as shown in Fig. 3. The fluorescent beads act as a point source which are employed to determine flow-variant detection PSF, and later on used to deconvolve the images at specific flow-rate. Three types of chip geometries are employed in the present study, one with a straight single channel and the others with two and four channels. As can be inferred from beads flow data (see, Fig. 3A), the count follows a linear increase with flow, whereas the same is not true for cells. This is due to many reasons including, cell accumulation at the corners of channels and adhesive nature of the cell that tend to make cluster. In addition, low cell count can be attributed to its relatively large mass, apart from other reasons. It may be noted that we have used cell culture medium as the specimen flowing fluid for the present study, and no additional chemicals (such as, Tripsine) are used in order to prevent clustering. Numerically, we got maximum count of 1421 cells per minute at 2000 *nl/min* when compared to > 2000 beads per minute. This is fantastic considering high resolution (organelle-level) volume imaging capability of SMMIC system.

**FIG. 3:**
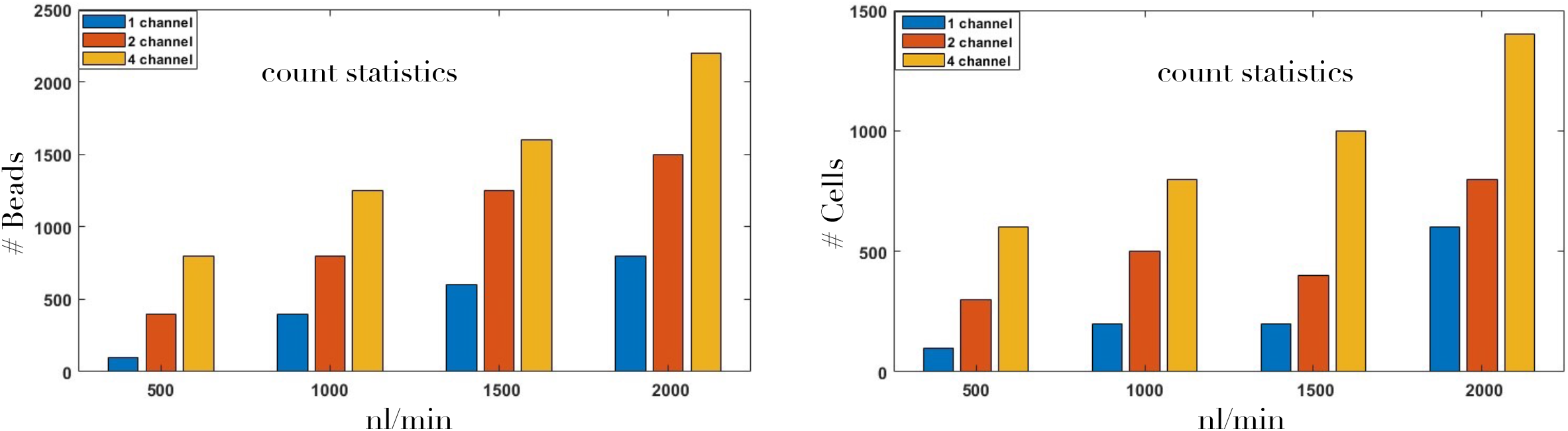
Flow statistics. (A) The number of beads as counted by the SMMIC system at varying flow-rates (500 - 2000 nl/min.). (B) The count of HeLa cells as they flow through the Y-type multichannel system transected by VAMSA PSF.

The quality of sectional images and the reconstructed volume are central to the newly developed SMMIC system. Fig. 4 shows simultaneous interrogation capability of the system in all the 4 channels along with the ability to reconstruct cell volume. The sectional images of the HeLa cells (spherical in shape during flow) displaying the distribution of organelle (mitochondria) are recorded by the sCMOS camera during flow (at 1000 *nl/min*). Five high quality planes of a sample HeLa cell (cell #1-5) are captured, which are then deconvolved using flow-variant PSF and stacked together to reconstruct the cell volume. The distribution of mitochondria inside a live HeLa cell can be ascertained from the volume images. The corresponding raw data of the cells passing through all the 4 channels can be found in **Supplementary Video 1**. Overall, this demonstrates the capability of newly developed SMMIC system for rapid interrogation of intracellular components with an organelle-level resolution at high throughput.

**FIG. 4:**
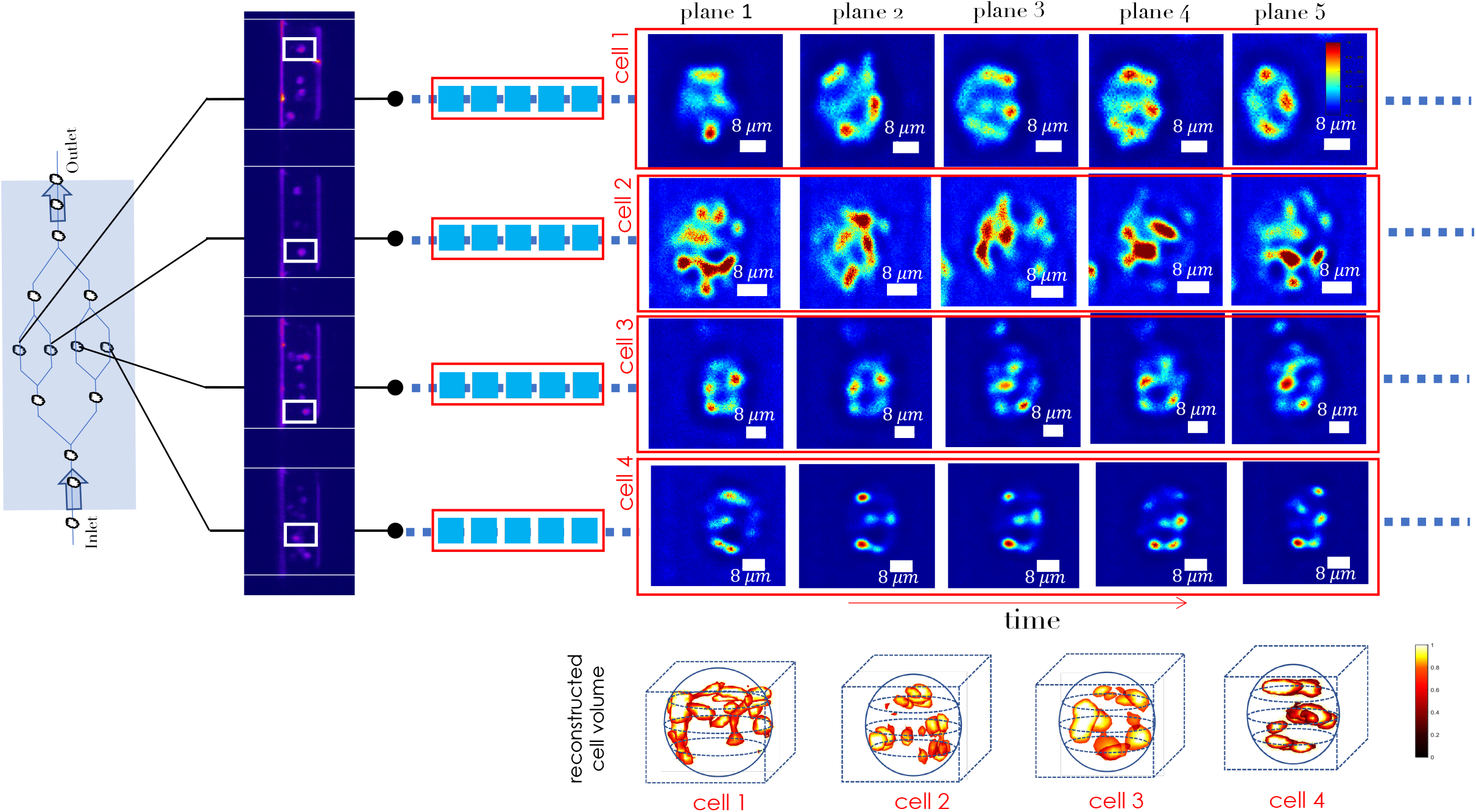
A snap-short of fluorescently-labelled cells flowing through the microfluidic channels as captured by the sensitive sCMOS camera. Several planes of the HeLa cells (indicated by white box) simultaneously passing through channels in the microfluidic chip are shown. The corresponding cell volumes reconstructed from the sectional images are also displayed. The raw experimental data is shown in the **Supplementary Video 1**.

The developed SMMIC system is compared with state-of-the-art point-illumination based imaging cytometry system and confocal microscopy, both in terms of performance and estimation of biophysical parameters as shown in Table 1. The study confirms that although traditional imaging cytometry is 10 times faster but SMMIC system has the distinct ability to reconstruct 3D volume during flow. The system facilitates organelle count in a cell volume which is largely due to its near diffraction-limited resolution. Moreover, the proposed light sheet powered system outperforms traditional imaging cytometry in terms of signal to background ratio (SBR). With the available computing power, the system is capable of reconstructing ∼ 121 cell volumes per minute (at 1000 *nl/min*). All these advantages makes SMMIC unique as far as precision is concerned (availability of multiple light sheets, and non-requirement of hydrodynamic focusing), portable (due to microfluidic chip based specimen flow platform) and viable as future commercial product (with the availability of additional features related to cell biophysical parameters and cell volume).

**TABLE I:** Key biophysical parameters (cell count, organelle count, # reconstructed 3D cell / cell volumes) and its performance comparison with existing cytometry systems.

| Technique | #cell count | #cell vol./min. | organelle count | resolution | SBR (dB) |
| --- | --- | --- | --- | --- | --- |
| Confocal | NA | NA | NA | organelle | 4.0 [39] |
| Traditional | > 10k | — | not-possible | cellular [40] | — |
| SMMIC (Raw) | > 1k | ~ 121 | possible | organelle | 1.67 |
| SMMIC (Deconvolved) | > 1k | ~ 121 | possible | organelle | 4.80 |

## Conclusion & Discussion

A newly developed imaging cytometry system (SMMIC) is demonstrated and successfully tested on cancerous HeLa cell samples. The system is realized primarily based on the generation of diffraction-limited VAMSA PSF aided by a specific configuration of transmission grating, beam-expander, cylindrical lens and high NA objective. The selectivity of the number of vertical light sheet and the adaptive change of inter-sheet distance makes the system versatile, and ensures optimal performance in terms of selective illumination of microfluidic channels. In addition, a flow variant detection PSF ensures high quality volume reconstruction with an increase in both SBR and resolution. The system allows rapid volume visualization at high throughput (> 1400 cells per minute) which is a distinct feature of SMMIC system. Studies on cancerous HeLa cells reveal healthy count rate and instant volume reconstruction (121 per minute) with an organelle-level resolution and SBR approaching confocal microscopy. Moreover, SMMIC has the potential to carry out a variety of studies ranging from drug-screening investigation to clinical health-care (e.g., Hematology, Pathology and Urology). We anticipate the system to vastly advance the field of translational medicine and clinical diagnosis.

## Acknowledgements

The authors acknowledge financial support from parent institute (Indian Institute of Science, Bangalore, India). PPM conceived the idea. PJ, PK, AS, and PPM carried out the experiment. PJ and AS prepared the samples. PPM wrote the paper by taking inputs from all the authors.

## Data Availability

The data that support the findings of this study are available from the corresponding author upon request.

